# Lesion Attributes Segmentation for Melanoma Detection with Deep Learning

**DOI:** 10.1101/381855

**Authors:** Eric Z. Chen, Xu Dong, Junyan Wu, Hongda Jiang, Xiaoxiao Li, Ruichen Rong

## Abstract

Melanoma is the most deadly form of skin cancer world-wide. Many efforts have been made for early detection of melanoma. The International Skin Imaging Collaboration (ISIC) hosted the 2018 Challenges to improve the diagnosis of melanoma based on dermoscopic images. In this paper, we describe our solution for the task 2 of ISIC 2018 Challenges. We present a multi-task U-Net model to automatically detect lesion attributes of melanoma. Our multi-task U-Net deep learning model achieves a Jaccard index of 0.433 on official test data, which ranks the 5th place on the final leaderboard.

## 1. INTRODUCTION

Skin cancer is one of the most common cancers worldwide and more than one million skin cancers have been diagnosed in the United States each year. Melanoma is the most dangerous form of skin cancer, which causes over 9,000 deaths each year [1]. Melanoma in the late stage can often spread to other parts of the body and thus it is difficult to treat and usually can be fatal. However, melanoma in the early stage is treatable and the majority can be cured. Many efforts have been made to detect melanoma in the early stage. Dermoscopy is one noninvasive method commonly used in the healthcare to examine pigmented skin lesion. It can generate high-resolution images of the lesion regions on the skin. To diagnose melanoma, it still requires the dermatologist to evaluate the images based on several skin lesion patterns. The common dermoscopic attributes are pigment network, negative network, streaks, milia-like cysts, and globules [2]. Automatic detection of those skin lesion attributes can be a tremendous help for early melanoma diagnosis.

Towards this goal, the International Skin Imaging Collaboration (ISIC) hosted the 2018 Challenges to diagnose melanoma automatically based on dermoscopic images. The 2018 challenges include three tasks. The first task is to predict lesion segmentation boundaries in dermoscopic images. The second task is to predict the locations of five skin lesion patterns (i.e., dermoscopic attributes) in dermoscopic images, which are pigment network, negative network, streaks, milia-like cysts, and globules. The third task is to predict seven disease categories in dermoscopic images, which includes melanoma, melanocytic nevus, basal cell carcinoma, actinic keratosis / Bowen’s disease, benign keratosis, dermatofibroma and vascular lesion. The three tasks mimic the steps for lesion analysis performed by dermatologists in the clinic settings. From the previous years’ competitions, it has been reported that far less participation in the second task of lesion pattern prediction than in other tasks [3]. It seems that the second task is the most difficult one and this motivated us to focus on this problem instead of the other two.

Deep learning, especially convolutional neural network (CNN), has been widely applied to solve many problems in computer vision. Various CNN based models have been developed for object classification and detection, such as VGG [4], ResNet [5], Inception [6], DenseNet [7], and more [8]. For image segmentation, FCN [9], U-Net [10], Mask R-CNN [11] are some of the popular deep learning models. Many deep learning based approaches have been applied for melanoma detection [2]. However, most of those researches applied deep learning models to either classify skin diseases [12, 13] or segment whole lesion regions [14], which are corresponding to the task 1 and task 3 in ISIC 2018 Challenges. To our knowledge, currently there is no deep learning based approach has been applied to segment specific lesion attributes as in the task 2 of ISIC 2018 Challenges. Note that in the previous ISIC Challenges, task 2 was based on the segmentation of superpixels [3].

In this paper, we describe our solution for the task 2 of ISIC 2018 Challenges. We present a multi-task U-Net model to automatically detect lesion attributes of melanoma. In the U-Net model, we replace the encoder part of the U-Net with a pretrained VGG16 network [15]. In the middle layer and the last layer of the U-Net, we add two classification heads to classify the empty masks versus the non-empty masks. Our multi-task U-Net model achieves a Jaccard index of 0.433 on official test data, which ranks the 5th place on the final leaderboard. The code for our solutions is publicly available at https://github.com/chvlyl/ISIC2018.

## 2. METHODS

### 2.1. Data

We downloaded the task 2 dataset from ISIC 2018 Challenges website [16]. For each skin image, this task is aimed at detecting five lesion attributes (pigment network, negative network, streaks, milia-like cysts, and globules). The dataset consists of 2,594 images and 12,970 ground truth masks (five masks for each skin image) as training data. The skin images are RGB images in JPG format and the masks are grayscale images in PNG format. The five masks are corresponding to five lesion attributes in each skin image. Note that some of the attributes may not present in skin images, therefore the corresponding masks are empty. There are 100 images provided as validation data and 1,000 images as test data. No ground truth masks are provided for validation data and test data during the competition.

We noticed that not all attributes present in each skin image (Figure 1), which motivated us to propose the following multi-task U-Net model.

**Fig 1.**
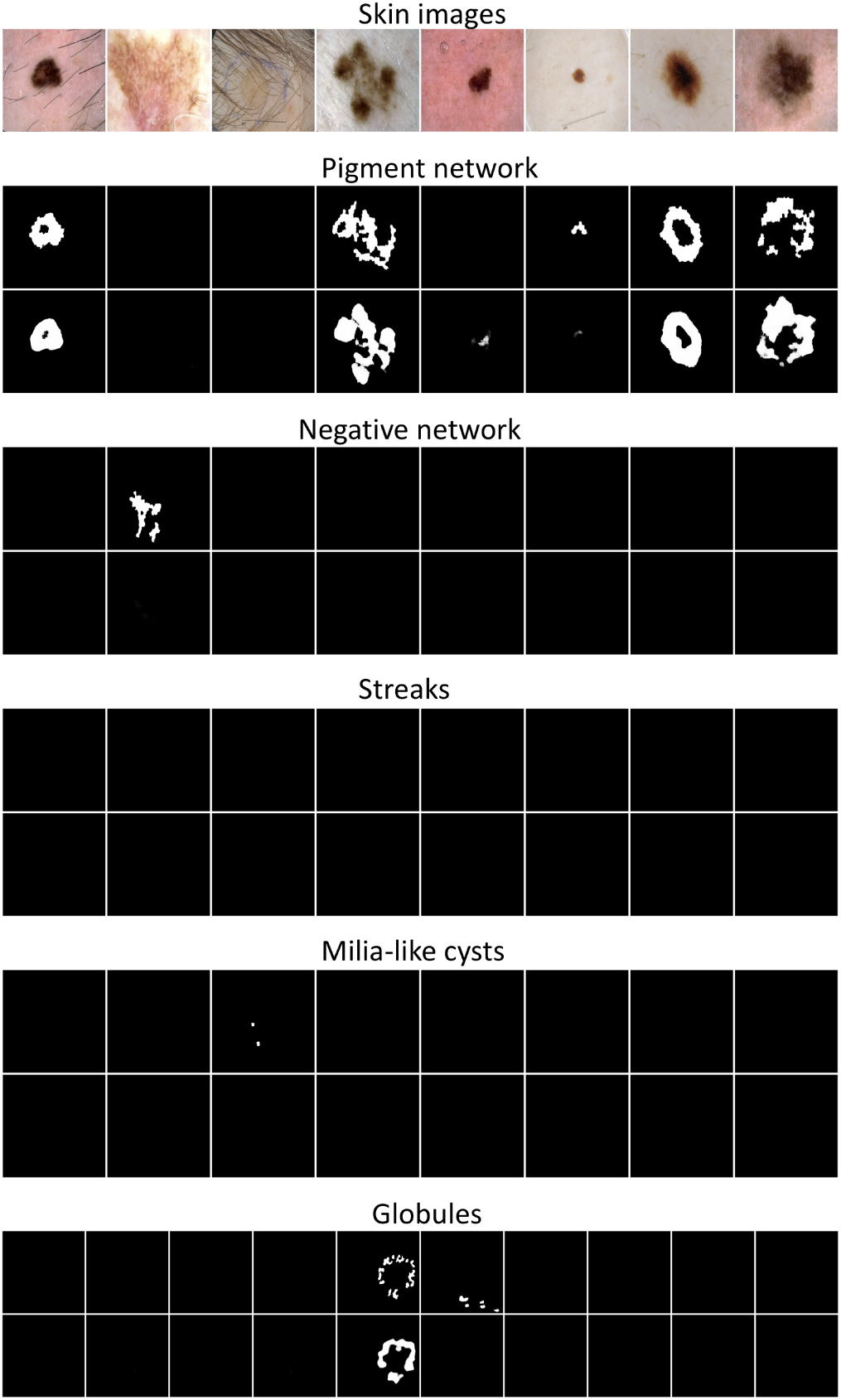
Example of skin images and five corresponding lesion attributes. For each attribute, the top row is the ground truth masks and the bottom row is the predicted masks from multitask U-Net model.

### 2.2. Multi-task U-Net model

Based on the original U-Net [10], we replace the encoder part with a pretrained VGG16 [17]. The pretrained weights are based on ImageNet data. In the skip connections, the feature maps from the encoder part are concatenated with the corresponding feature maps in the decoder part. The output of the U-Net is a five channel mask. We define the loss function for pixel wise segmentation as a combination of binary cross entropy loss L and Jaccard index J [15].

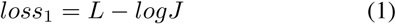

We observed that for each skin images, not all lesion attributes present in the masks. Therefore, we also add two classification heads to classify the empty masks versus the non-empty masks. Since the last layer of our U-Net is a convolutional layer and the output is five feature maps, we add a global max pooling layer to this last convolutional layer and use this to classify the empty and non-empty masks for five attributes. This classification head will act as a regularizer. Since if the truth mask is empty and our network predicts a non-empty mask, after global max pooling, the classification loss will be large. Therefore, this classification head will force the network to predict an empty mask if the true mask is empty. We also adopt the similar idea in Inception [6] and add another classification head to the middle layer with a 1 × 1 convolutional layer and a global average pooling layer. The network structure is shown in Figure 2.

**Fig 2.**
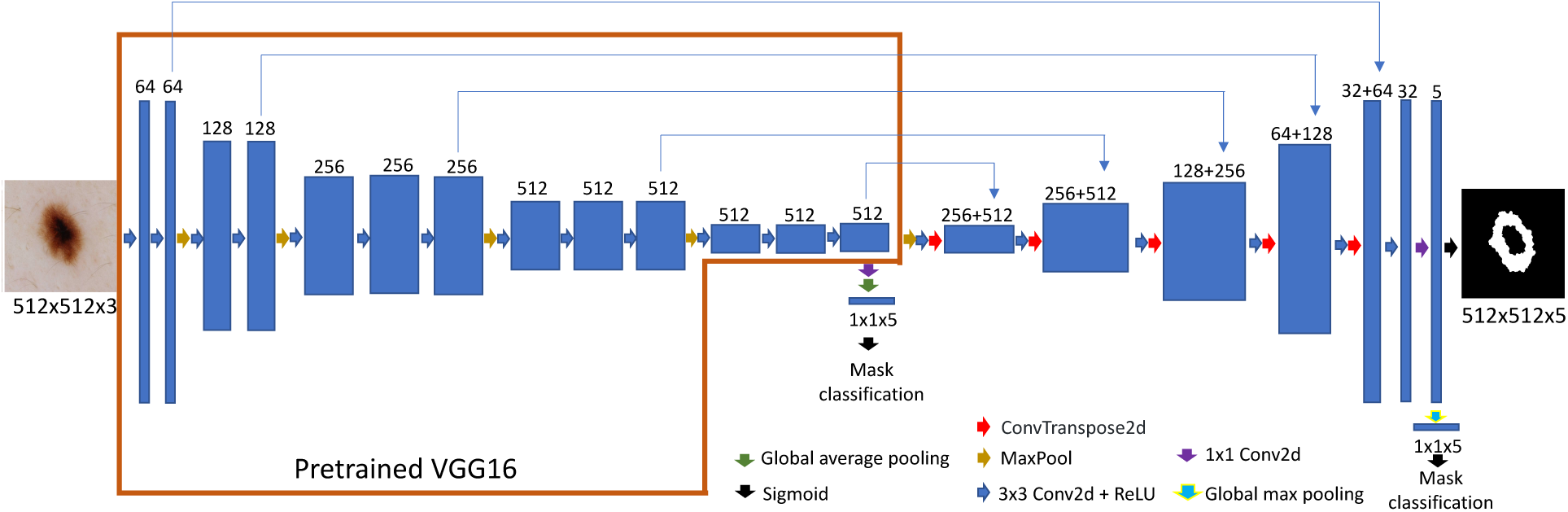
Network structure using U-Net architecture. The encoder part of the U-Net is replaced with a pretrained VGG16 network [15]. We also add two classification heads to the network, where one is added to the middle layer with 1 × 1 convolution and global average pooling layers and the other is added to the last layer with a global max pooling layer.

The final loss is a combination of pixel wise segmentation loss *loss*_1_, classification loss using the middle layer *loss*_2_ and classification loss using the last layer *loss*_3_. We used binary cross entropy for *loss*_2_ and *loss*_3_. We assigned weights 0.5 to the last two losses.

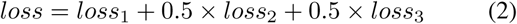

### 2.3. Model training

The raw skin images have various sizes, from 540 *×* 722 to 4499 *×* 6748. We first resize all the skin images and corresponding masks to 512 *×* 512. Then for each skin image, we concatenate the five corresponding grayscale masks into a five-channel mask. Each channel is a 512 *×* 512 mask. Both skin images and mask images are divided by 255 to scale the values in [0,1].

We first freeze all the encoder layers and only train the decoder layers for 50 epochs. Then we train the whole network for another 250 epochs. We use a learning rate of 0.01 with Adam optimizer and then decrease it by 0.8 when validation loss does not improve for 5 epochs during training. During training, we augment the skin images and masks by random flipping, rotation, scaling as well as randomly adjusting brightness and saturation. We use cutoff 0.3 to transform the predicted probability into the binary mask. The whole analysis is implemented in Pytorch.

## 3. RESULTS

Figure 1 shows an example of several skin images and their corresponding lesion attributes (for each attribute, top rows show the ground truth masks). We noticed that not all attributes present in each skin image. That is, some of the masks are empty for certain skin images. Table 1 shows the summary of non-empty masks for each lesion attribute. 58.7%, 26.3% and 23.2% of the skin images have non-empty masks for Pigment network, milia-like cysts, and globules, respectively. While only 7.3% and 2.9% of the skin images have non-empty masks for negative network and streaks, respectively. Since segmentation is essentially a pixel-wise classification problem, a large number of empty masks increase the number of samples in negative class. In the U-Net model, we add two classification heads to classify the empty versus nonempty masks. Figure 1 shows the predicted masks from the multi-task U-Net model (for each attribute, the bottom rows show the predicted masks). The Jaccard index is 0.477 on the official validation dataset. The final Jaccard index on the official test data is 0.433, which ranks the 5th place on the final leaderboard.

**Table 1.** Summary of non-empty masks in the training data

| Attributes | Number of non-empty masks | Percentage |
| --- | --- | --- |
| Pigment network | 1522 | 58.7% |
| Negative network | 189 | 7.3% |
| Streaks | 100 | 2.9% |
| Milia-like cysts | 681 | 26.3% |
| Globules | 602 | 23.2% |
| Total skin images | 2594 | 100% |

## 4. DISCUSSION

In this paper, we propose a multi-task U-Net model to detect five attributes of skin lesions. We replace the encoder part of the U-Net with a pretrained VGG16 network and add two classification heads to the U-Net. Our submission ranks the 5th place on the final leaderboard of ISIC 2018 task 2.

For our U-Net based model, we tried to add the batch normalize layers to the encoder and decoder parts of the network. However, this does not improve the model performance compared to the model without batch normalization layers. We also tried to replace the encoder part of the U-Net network with pretrained DenseNet. However, the GPU memory usage was too hight and we could only fit one image per patch. The training took a long time and the final result was worse than that with VGG16 as the encoder.

We also noticed that the five lesion attributes are highly imbalanced in the training data. For example, 58.7% of the skin images have the pigment network attribute but only 2.9% of skin images have the negative network attribute. Classification of imbalanced data is often difficult. One potential solution is to assign different weighs to each attribute during training. However, we spent some time exploring different weight configurations but it seems to us there was no significant improvement.

Mask R-CNN has been recognized as the state-of-art neutral network architecture for instance segmentation. We also applied the Mask R-CNN architecture for the skin attribute detection task in the challenge. The best Jaccard index on the official validation data is 0.233 and we did not submit our test result for the Mask R-CNN model.

